# Emerging Pathogens in Urinary Tract Infections: Virulence and Phenotypic Characterization of *Pseudomonas aeruginosa* strains

**DOI:** 10.64898/2026.01.08.698422

**Authors:** Rachel C. Fleck, Olena Gorodyna, Hao Zhou, Hannah Bryant, Mihaela Gadjeva, Allyson E. Shea

## Abstract

Urinary tract infections (UTIs) affect a broad patient population and inflict a substantial financial burden on the U.S. healthcare system. While uropathogenic *Escherichia coli* (UPEC) causes the majority of cases, other pathogens are emerging. Analysis of patient data from our healthcare system in the Gulf Coast region of Alabama revealed that *Pseudomonas aeruginosa* accounted for 4.0% of UTI cases, roughly double the national average, prompting further investigation into this historically understudied uropathogen. Here, we performed whole-genome sequencing and phenotypic assays on 55 urinary *P. aeruginosa* isolates to identify key drivers of pathogenicity in the context of UTI. Multilocus sequence typing identified 19 novel sequence types, underscoring the uncharacterized diversity of urinary *P. aeruginosa* isolates. Serotype O6 was most common and enriched in patients with indwelling catheters, whereas O4 was linked to diabetes mellitus. Antibiotic susceptibility testing (AST) revealed high levofloxacin resistance (30.9%), with 23.6% multidrug-resistant (MDR) and 9.1% extensively drug-resistant (XDR) isolates. Resistance patterns correlated with demographics, including significantly higher meropenem and aztreonam resistance in isolates from African American patients. Phenotypic assays of growth, motility, and biofilm formation revealed negative correlations between antibiotic resistance and virulence. Specific virulence genes predicted enhanced iron acquisition, hemolysis, and colonization potential. Notably, motility and exotoxin profiles emerged as strong predictors of *P. aeruginosa* ascension in a murine UTI model. Together, these findings provide new biological and clinical insight into *P. aeruginosa* as a uropathogen and emphasize the need for continued research.

**IMPORTANCE:** *Pseudomonas aeruginosa* is an emerging but understudied pathogen in urinary tract infections (UTIs). Given its resilience, adaptability, and the growing threat of multidrug resistance, *P. aeruginosa* remains a significant challenge in clinical microbiology and infection control. Our data reveal an increased prevalence of *P. aeruginosa* in our local patient population. In this study, we examined both genotypic and phenotypic traits of clinical isolates and correlated them with colonization in murine models and extensive patient metadata. We identified strong associations between antibiotic resistance patterns and patient demographics. Novel sequence type strains were linked to motility phenotypes *in vitro*. Additionally, specific flagellar alleles were associated with enhanced murine kidney colonization and recurrent UTIs in patients. These findings provide new insight into the evolutionary adaptations that contribute to *P. aeruginosa* uropathogenicity and support a more nuanced understanding of its clinical significance.

## INTRODUCTION

*Pseudomonas aeruginosa* is a Gram-negative, opportunistic pathogen best known for causing severe lung and wound infections, yet it can also pose significant challenges in the urinary tract^1^. Although urinary tract infections (UTIs) are predominantly caused by uropathogenic *Escherichia coli* (UPEC), *P. aeruginosa* accounts for up to 10% of catheter-associated urinary tract infections (CAUTIs) and 16% of UTIs in intensive care units (ICUs)^2–4^. The prevalence of *P. aeruginosa* in hospital-acquired infections is driven by innate resistance to many antibiotics, adaptability to diverse environments, and numerous virulence factors. However, in contrast to extensive studies in lung and wound infection models, the characteristics that allow *P. aeruginosa* to successfully colonize the urinary tract remain poorly understood.

A defining feature of *P. aeruginosa* is its extensive intrinsic and acquired antibiotic resistance mechanisms. Its large genome encodes multiple efflux pumps, β-lactamases, and aminoglycoside-modifying enzymes, conferring innate resistance to a broad range of antimicrobial classes^5^. *Pseudomonas* clinical isolates are often multi-drug resistant (MDR), defined as resistance to ≥1 agent in ≥3 antimicrobial classes, or extensively drug-resistant (XDR), defined as resistance to all but one or two classes^6^. Approximately 29% of complicated UTIs (cUTIs) caused by *P. aeruginosa* are due to MDR strains, making treatment especially challenging^7^. Because of this, the Centers for Disease Control and Prevention (CDC) has classified MDR *P. aeruginosa* as a serious public health threat^8^.

In addition to antimicrobial resistance, *P. aeruginosa* employs a broad range of virulence mechanisms. Biofilm formation, particularly on urinary catheters, protects bacteria from host defenses as well as antibiotic penetration and relies on flagellar motility genes (type A *flaA* and type B *fliC*) and pili genes such as *pilA*^1,9^. Quorum sensing (QS) further coordinates biofilm formation and persistence by linking environmental sensing to virulence expression via key transcriptional regulators such as *lasR* and *rhlR*^10^. Given the rise of antibiotic resistance, these virulence mechanisms have emerged as targets of antivirulence therapies^11,12^. Indeed, experimental disruption of QS pathways has been shown to significantly reduce virulence in the urinary tract^13,14^ , highlighting the importance of researching these mechanisms to better understand and treat *P. aeruginosa* UTIs.

Another critical virulence feature is the secretion of exotoxins, particularly via the type III secretion system (T3SS). The T3SS injects cytotoxins such as ExoS, ExoT, ExoU, and ExoY directly into host cells, disrupting cytoskeletal integrity, interfering with immune signaling, and promoting cell death^15–18^. Interestingly, the T3SS was shown to be essential for establishing acute infection but not required for persistence in chronic infection in a murine model of CAUTI^19^. Among these four exotoxins, ExoU and ExoS are considered most clinically relevant and are thought to be mutually exclusive in *P. aeruginosa* strains^20,21^. ExoU is particularly associated with virulence, severe cytotoxicity, and poor clinical outcomes in the context of pneumonia^22,23^. In addition to the T3SS, the type II secretion system (T2SS) also contributes to *P. aeruginosa* virulence via secretion of exotoxin A (ToxA), a highly cytotoxic protein strongly linked to pneumonia and chronic lung infections in cystic fibrosis patients^24–26^. Expression of exotoxin A is directly regulated by the iron-scavenging siderophore pyroverdine, partially encoded by *pvdA*, which is essential for growth in iron-limited environments such as urine^27–29^. No single virulence factor alone drives *P. aeruginosa* pathogenesis, but rather it is the combination of these factors that enables successful infection of the urinary tract.

In addition to virulence factors, variability in *P. aeruginosa* serotypes can also influence infection severity and treatment outcomes. Among the 20 known *P. aeruginosa* O-antigen serotypes, O6 and O11 have been associated with increased mortality in patients with pneumonia^30^. Isolates of serotype O11 often harbor *exoU* and cause significant epithelial damage and have also been associated with multidrug resistance (MDR)^31,32^. Given their pathogenic potential, recent studies have investigated O-antigens as vaccine therapeutic targets, underscoring the importance of characterizing O-antigen serotypes in clinical *P. aeruginosa* isolates^33^. Patient-related factors also play a critical role in UTI susceptibility and recurrence. Several factors are known to increase the risk of *P. aeruginosa* UTIs, including indwelling urinary catheters (IDCs), immunosuppressive therapies, diabetes mellitus (DM), and prolonged hospitalization^7^. However, the roles of other host factors in *P. aeruginosa* infection and virulence in the urinary tract remain relatively understudied, highlighting the need for integrative studies that investigate both pathogen diversity and patient characteristics in the context of UTI.

Here, we phenotypically and genotypically assessed 55 *P. aeruginosa* clinical urinary isolates from a diverse patient population. We performed bacterial growth, biofilm formation, iron acquisition, cytotoxicity, and motility assays to directly assess *P. aeruginosa* virulence and persistence *in vitro*. We also conducted whole-genome sequencing (WGS) to determine the presence of important virulence and antibiotic resistance genes as well as characterize the multi-locus sequence type (MLST) and O-antigen serotype of each strain. With these results and abundant patient metadata, we correlated phenotype, genotype, and patient variables to uncover drivers of *P. aeruginosa* pathogenicity in UTI, validated by our *in vivo* murine UTI model. Understanding the interplay between these factors is crucial for developing effective therapeutic and UTI preventive strategies.

## RESULTS

### Emergence of novel genotypes in uropathogenic *P. aeruginosa* strains

*Pseudomonas aeruginosa* has been extensively studied in the context of lung and wound infections but relatively understudied as a urinary tract infection (UTI) pathogen. Nationwide, *P. aeruginosa* accounts for only 1% of uncomplicated UTIs and 2% of complicated UTIs^3^; however, we report a prevalence of 4.03% in our local healthcare system (**Fig. S1**). This discrepancy prompted us to investigate the genotypic and phenotypic characteristics of *P. aeruginosa* strains in our region and to explore their associations with patient variables to better understand clinical risk. Fifty-five *P. aeruginosa* clinical isolates were obtained from urine samples of patients within the University of South Alabama healthcare system, representing a diverse cohort with various comorbidities and known UTI risk factors (**Table 1**). Three reference strains were included in this study for comparison: PAO1, PA103, and PA27853, isolated from wound, lung, and bloodstream infections, respectively^34–37^. Indeed, there is no available *P. aeruginosa* urinary type strain for comparison.

**Table 1:** Patient demographics and comorbidities

| Patient Information |  |
| --- | --- |
| **Age** |  |
| Median [Min, Max] | 71 [0.33, 92] |
| **Sex** |  |
| Male | 26 (47.3%) |
| Female | 29 (52.7%) |
| **Race** |  |
| African American (AA) | 18 (32.7%) |
| White | 37 (67.3%) |
| **Ethnicity** |  |
| Hispanic | 2 (3.6%) |
| Not Hispanic | 53 (96.4%) |
| **BMI** |  |
| Median [Min, Max] | 27.3 [17, 65] |
| Missing | 9 (16.4%) |
| **Co-Infection** |  |
| Yes | 5 (9.1%) |
| No | 50 (90.9%) |
| **Comorbidities** |  |
| Sepsis | 7 (12.7%) |
| Diabetes mellitus (DM) | 22 (40.0%) |
| Reccurent UTI (rUTI) | 41 (74.5%) |
| Indwelling catheter (IDC) | 23 (41.8%) |

Whole-genome sequencing (WGS) was performed on all clinical isolates, and the genomes were assembled and annotated using BV-BRC^38^. A phylogenetic tree constructed from single nucleotide polymorphisms (SNPs) revealed multiple distinct clades, suggesting a high degree of genomic diversity among clinical isolates (**Fig. 1A**). Multilocus sequence typing (MLST) via PubMLST^39^ identified 19 novel sequence types (STs) (**Table 2**), further demonstrating the lack of genotypic characterization of uropathogenic *P. aeruginosa*. O-antigen serotyping was performed with the PAst *in silico* online tool^40^, and a total of 8 distinct serotypes were identified in our cohort, with O6 being the most prevalent (38.18%) (**Fig. 1B**). The O6 serotype was enriched in patients with indwelling catheters (IDCs) (**Fig. 1C**), while O4 strains were more common in patients with diabetes mellitus (DM) (**Fig. 1D**). These findings demonstrate previously uncharacterized diversity in urinary isolates, motivating further investigation into how this genetic variation shapes antibiotic resistance and virulence phenotypes.

**Figure 1.**
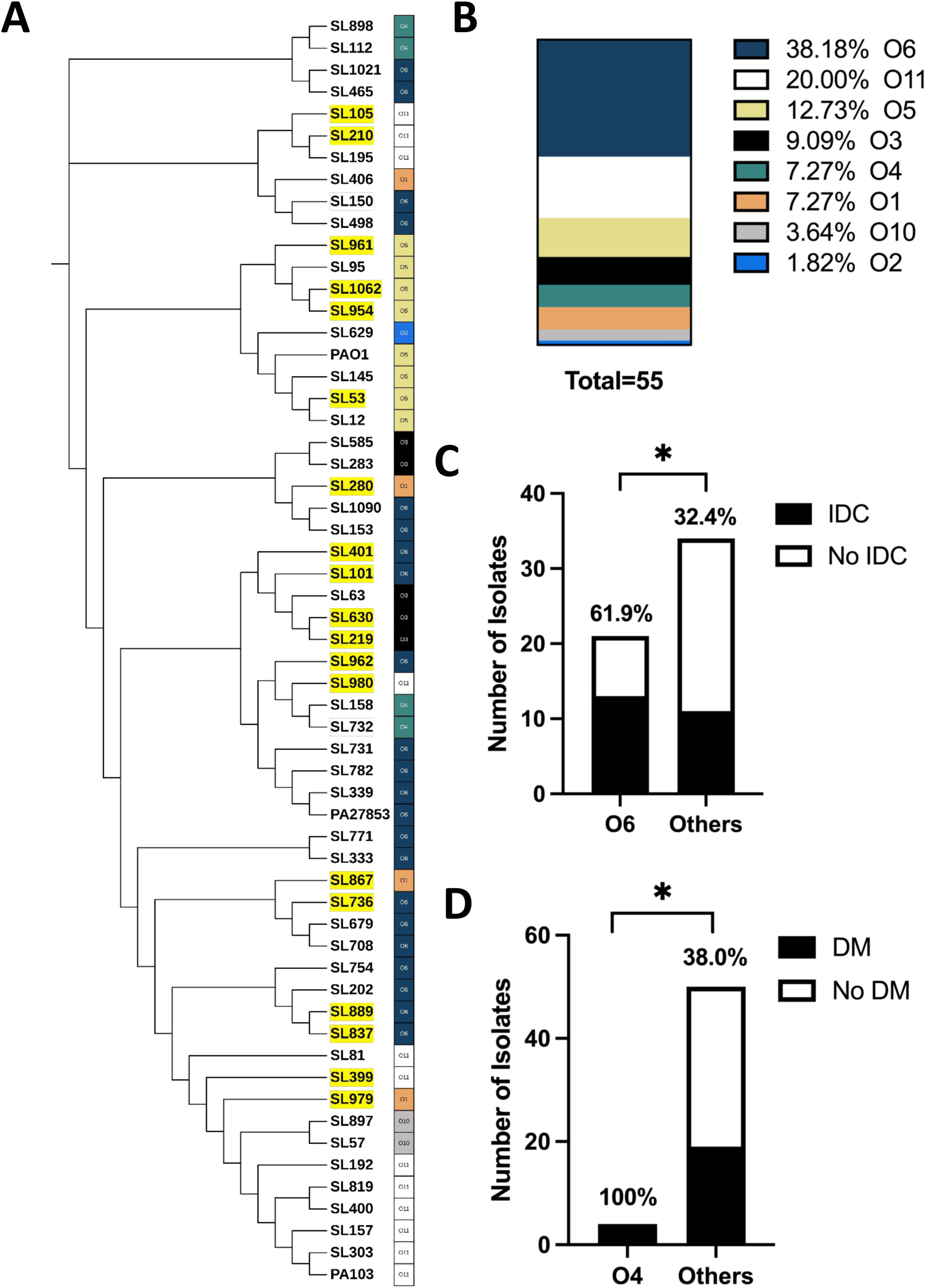
**Genotypic and phylogenetic characterization of 55 *Pseudomonas aeruginosa* UTI clinical isolates**. **(A)** Phylogenetic tree of 55 *P. aeruginosa* clinical isolates and type strains PAO1, PA103, and PA2785316 with novel ST types (highlighted in yellow) and serotypes indicated. **(B)** Serotype prevalence in the clinical isolates (n=55) determined by PAst. Bar graphs indicate the number of isolates from **(C)** patients with indwelling catheters (IDCs) for serotype O6 versus all other serotypes and **(D)** patients with diabetes mellitus (DM) for serotype O4 versus all others. Fisher’s exact test was used to determine significance between groups (*P<0.05).

**Table 2:** MLST profiles of clinical *P. aeruginosa* urinary isolates.

| Strain | Sequence Type | Allele |  |  |  |  |  |  |
| --- | --- | --- | --- | --- | --- | --- | --- | --- |
|  |  | *acsA* | *aroE* | *guaA* | *mutL* | *nuoD* | *ppsA* | *trpE* |
| SL12 | 244 | 17 | 5 | 12 | 3 | 14 | 4 | 7 |
| SL53 | Novel | 17 | 5 | 94* | 3 | 14 | 4 | 7 |
| SL57 | 1197 | 5 | 8 | 119 | 6 | 12 | 6 | 3 |
| SL63 | 274 | 23 | 5 | 11 | 7 | 1 | 12 | 7 |
| SL81 | 2629 | 28 | 20 | 94 | 3 | 1 | 4 | 10 |
| SL95 | 487 | 11 | 5 | 6 | 18 | 4 | 15 | 19 |
| SL101 | Novel | 148 | 3 | 1 | 3 | 1 | 124 | 181 |
| SL105 | Novel | 6 | 4 | 62 | 66 | 48 | 6 | 26 |
| SL112 | 2932 | 11 | 19 | 19 | 3 | 4 | 4 | 3 |
| SL145 | 244 | 17 | 5 | 12 | 3 | 14 | 4 | 7 |
| SL150 | 5080 | 39 | 5 | 36 | 72 | 1 | 42 | 19 |
| SL153 | 2680 | 6 | 130 | 1 | 5 | 4 | 4 | 17 |
| SL157 | 298 | 18 | 4 | 13 | 3 | 1 | 17 | 13 |
| SL158 | 485 | 11 | 76 | 5 | 3 | 61 | 14 | 3 |
| SL192 | 2555 | 13 | 4 | 9 | 3 | 1 | 6 | 9 |
| SL195 | 697 | 17 | 24 | 37 | 5 | 3 | 7 | 25 |
| SL202 | 233 | 16 | 5 | 30 | 11 | 4 | 31 | 41 |
| SL210 | Novel | 17 | 89* | 37 | 5 | 3 | 7 | 25 |
| SL219 | Novel | 125 | 89* | 11 | 7 | 1 | 12 | 7 |
| SL280 | Novel | 6 | 5 | 5 | 3 | 3 | 6 | 11* |
| SL283 | 1229 | 28 | 5 | 11 | 11 | 4 | 15 | 7 |
| SL303 | 298 | 18 | 4 | 13 | 3 | 1 | 17 | 13 |
| SL333 | 2652 | 40 | 5 | 30 | 3 | 3 | 4 | 14 |
| SL339 | 179 | 36 | 27 | 28 | 3 | 4 | 13 | 7 |
| SL399 | Novel | 15 | 17 | 94 | 11 | 4 | 4 | 10 |
| SL400 | 446 | 18 | 4 | 5 | 3 | 1 | 17 | 13 |
| SL401 | Novel | 28 | 3 | 11 | 3 | 85 | 42 | 91 |
| SL406 | 27 | 6 | 5 | 6 | 7 | 4 | 6 | 7 |
| SL465 | 2948 | 16 | 5 | 36 | 3 | 4 | 13 | 14 |
| SL498 | 2900 | 11 | 8 | 6 | 5 | 1 | 13 | 14 |
| SL585 | 198 | 11 | 5 | 11 | 11 | 3 | 27 | 7 |
| SL629 | 1667 | 17 | 143 | 6 | 1 | 4 | 7 | 9 |
| SL630 | Novel | 23 | 5 | 11 | 7 | 1 | 262* | 7 |
| SL679 | 633 | 125 | 5 | 6 | 3 | 4 | 13 | 23 |
| SL708 | 633 | 125 | 5 | 6 | 3 | 4 | 13 | 23 |
| SL731 | 2633 | 11 | 177 | 11 | 18 | 3 | 4 | 3 |
| SL732 | 1033 | 11 | 22 | 5 | 3 | 61 | 14 | 3 |
| SL736 | Novel | 11 | 5 | 6 | 3 | 3 | 7 | 47 |
| SL754 | 782 | 15 | 3 | 3 | 11 | 1 | 15 | 1 |
| SL771 | 390 | 39 | 5 | 1 | 3 | 4 | 46 | 56 |
| SL782 | 4759 | 11 | 5 | 11 | 3 | 73 | 4 | 14 |
| SL819 | 446 | 18 | 4 | 5 | 3 | 1 | 17 | 13 |
| SL837 | Novel | 113 | 5 | 30 | 11 | 4 | 31 | 7 |
| SL867 | Novel | 28 | 5 | 226 | 2 | 1 | 4 | 10 |
| SL889 | Novel | 113 | 5 | 30 | 11 | 4 | 31 | 7 |
| SL897 | 2406 | 29 | 8 | 36 | 67 | 1 | 6 | 3 |
| SL898 | 234 | 1 | 5 | 11 | 3 | 4 | 10 | 3 |
| SL954 | Novel | 17 | 20 | 6 | 34 | 3 | 6 | 19 |
| SL961 | Novel | 11 | 129 | 1 | 3 | 4 | 6 | 3 |
| SL962 | Novel | 17 | 57 | 7 | 3 | 4 | 4 | 1 |
| SL979 | Novel | 32 | 4 | 5 | 13 | 1 | 7* | 4 |
| SL980 | Novel | 6 | 5 | 122 | 3 | 4 | 6 | 7 |
| SL1021 | 505 | 6 | 20 | 1 | 11 | 4 | 4 | 2 |
| SL1062 | Novel | 17 | 20 | 6 | 34 | 3 | 6 | 19 |
| SL1090 | 1074 | 6 | 130 | 1 | 4 | 2 | 4 | 17 |
| PA01 | 549 | 7 | 5 | 12 | 3 | 4 | 1 | 7 |
| PA103 | 298 | 18 | 4 | 13 | 3 | 1 | 17 | 13 |
| PA2785316 | 155 | 28 | 5 | 36 | 3 | 3 | 13 | 7 |

\*imperfect match

### Patient demographics strongly associate with specific antibiotic resistance profiles in *P. aeruginosa* UTI strains

Antibiotic resistance is becoming an urgent global threat, and *P. aeruginosa* is a noted pathogen of high importance (ESKAPE pathogen) due to its innate and acquired resistance to many antibiotics^41^. First-line therapies often include the cephalosporins cefepime and ceftazidime, or the fluoroquinolones ciprofloxacin and levofloxacin, while meropenem is often reserved for highly resistant strains^42^. In our cohort, antimicrobial susceptibility testing (AST) revealed a wide range of resistance profiles to seven clinically relevant antibiotics (**Fig. 2A**). Levofloxacin resistance was the most common, with 36.4% (20/55) of isolates conferring resistance. Of the 55 strains, 13 (23.6%) were multidrug-resistant (MDR) and 5 (9.1%) were extensively drug-resistant (XDR). Isolates of the O11 serotype were 7.9 times more likely to be XDR (**Fig. S2**). Interestingly, resistance patterns varied by specific patient demographics and comorbidities. Isolates from African American patients were significantly more likely to be resistant to meropenem (**Fig. 2B**) and 14.2 times more likely to have intermediate aztreonam resistance (**Fig. 2C**) compared to those from White patients. Intermediate cefepime resistance was more common among strains isolated from polymicrobial infections than single causative agent cases (**Fig. 2D**), and aztreonam resistance positively correlated with the presence of novel sequence types (**Fig. 2E**). In addition, isolates from male patients harbored significantly more resistances per strain than those from female patients (**Fig. 2F**), with 50% of male-derived isolates and 24% of female-derived isolates having 1 or more resistance. To complement phenotypic testing, we also assessed antibiotic resistance genotypically using the Comprehensive Antibiotic Resistance Database (CARD)^43^ (**Fig. S3**) and discovered the presence of the MDR-associated gene *armR* was negatively correlated with patient BMI (**Fig. 2G, Fig. S4**). Together, these patterns point to complex drivers of resistance that span bacterial genotype, host demographics, and infection context.

**Figure 2.**
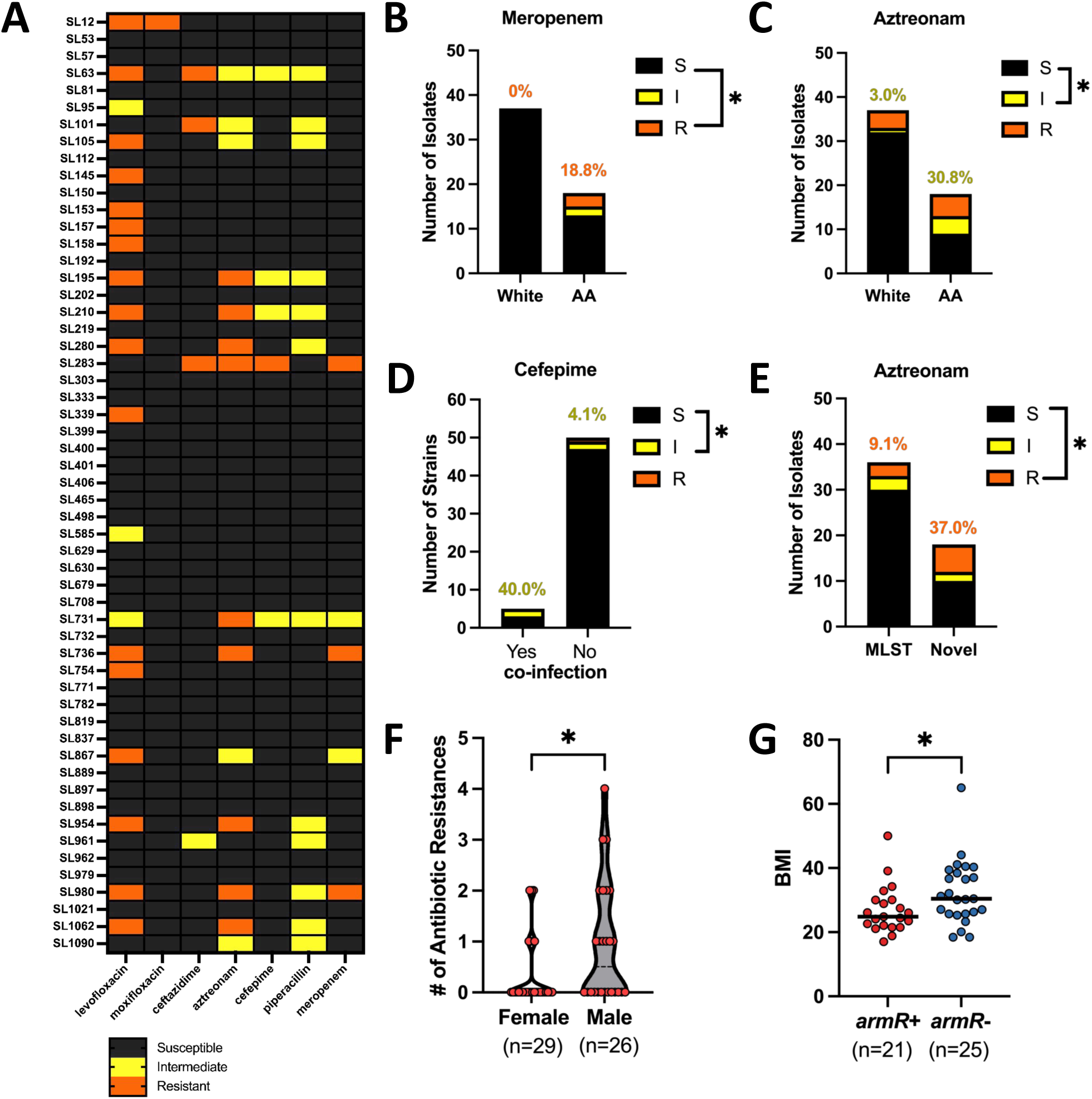
Antimicrobial resistance in *P. aeruginosa* isolates is associated with patient race, sex, and strain type. **(A)** Antibiotic susceptibility testing (AST) was performed at the University of South Alabama hospital, and strains were classified as susceptible (S, shown in black), intermediate (I, shown in yellow), or resistant (R, shown in orange) to 7 antibiotics based on the minimum inhibitory concentrations (MICs). **(B-C)** Bar graphs representing the median number of isolates from White and African American (AA) patients with **(B)** meropenem resistance and **(C)** aztreonam intermediate resistance. **(D)** Bar graph of cefepime intermediate susceptibility in isolates from polymicrobial vs monomicrobial UTIs. **(E)** Bar graph representing aztreonam susceptibility in isolates with established ST types (MLST) vs novel ST types (Novel). Fisher’s exact test was used to determine significance of panels B-E (*P<0.05). **(F)** Violin plot of the total number of antibiotic resistances (R or I) per strain in isolates from females and males. Each point represents a clinical isolate. **(G)** Plot of patient BMI of isolates with (red) and without (blue) the *armR* gene. Each dot represents a clinical isolate, and the median is indicated with a black line. Mann-Whitney U test was used for panels F-G (*P<0.05).

### Virulence-associated genes predict differential growth in human urine

To assess bacterial fitness under both nutrient-rich and host-relevant conditions, we quantified the growth of 55 *P. aeruginosa* clinical isolates and 3 reference strains in Luria Broth (LB) and filter-sterilized, pooled human urine. Many clinical isolates outperformed the reference strains in either medium, but none excelled in both LB and human urine, suggesting a metabolic trade-off (**Fig. 3A**). The presence of virulence genes was then determined by Basic Local Alignment Search Tool Protein (BLASTP), and these results were correlated with growth. Because gene detection was based on similarity to PAO1 alleles, some genes classified as “absent” may instead represent divergent urinary variants with <85% amino acid identity. Growth in human urine was enhanced 1.2-fold in isolates harboring AMR-associated genes *parS* and *armR* and 1.5-fold in isolates with the virulence gene *toxA* (**Fig. S5A**). Additionally, strains with the O5 serotype showed a 20.7% growth increase (**Fig. S5B**) while strains with serotype O11 exhibited a 28.9% growth decrease in human urine (**Fig. S6C**) compared to other serotypes. Strains isolated from polymicrobial UTIs showed a 36.6% increase in human urine growth (**Fig. 3B**) compared to isolates from single organism infections. In LB, strains with virulence factors *exoU* (**Fig. 3C**) and *aprA* (**Fig. 3D**) as well as those with pili-associated gene *pilA* (**Fig. 3E**) displayed significantly reduced growth. Interestingly, no patient-associated variables aside from co-infection correlated with pathogen growth (**Fig. S6**). Overall, these findings indicate that specific virulence factors not only affect pathogenic potential but may also shape metabolic adaptability in the urinary tract.

**Figure 3.**
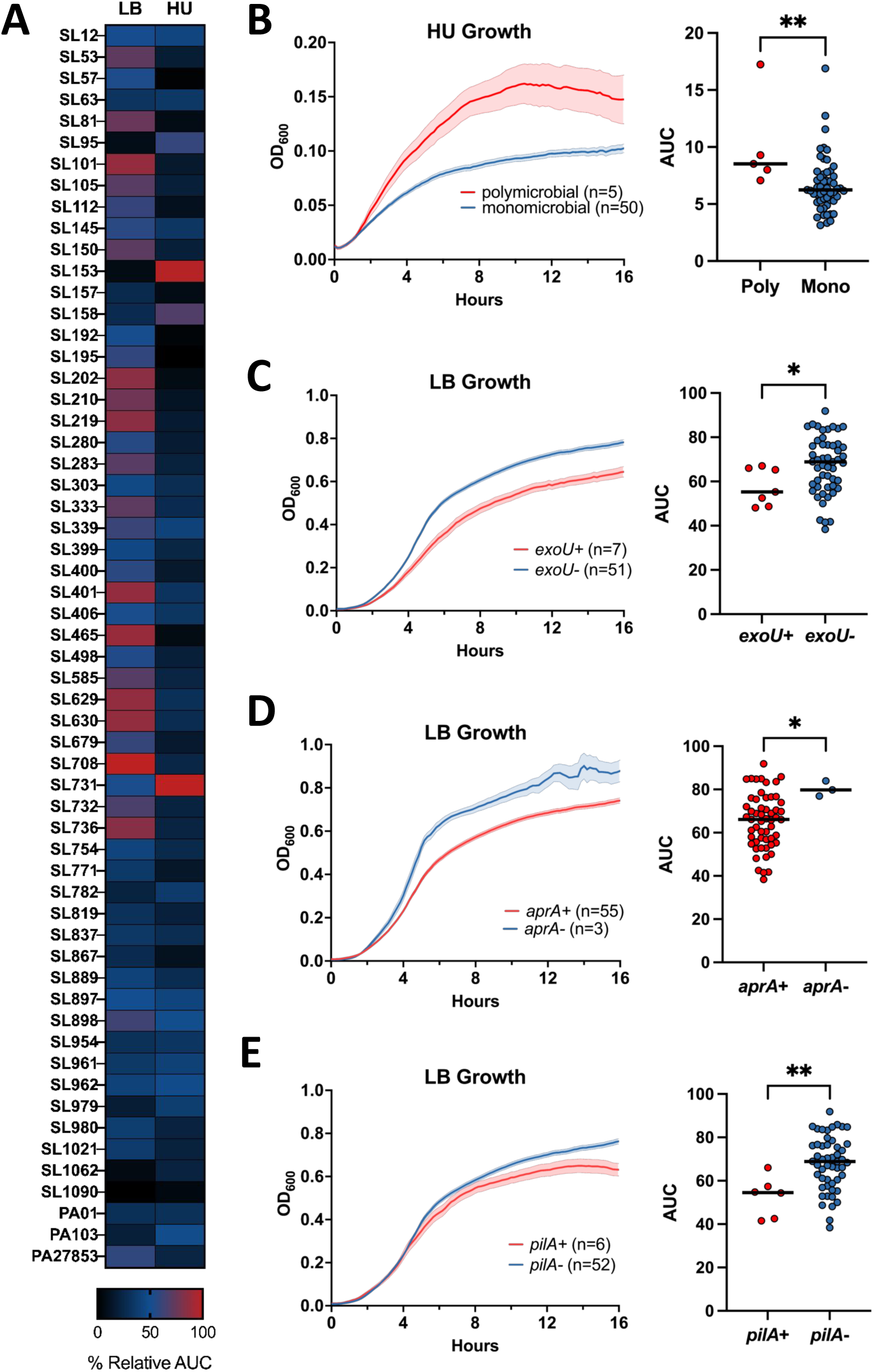
Virulence genes and polymicrobial infection influence growth of *P. aeruginosa*. **(A)** Growth of *P. aeruginosa* clinical isolates and type strains PAO1, PA103, and PA2785316 in LB Broth (LB) and filter-sterilized pooled human urine (HU) displayed in a heatmap of normalized area under the curve (AUC) (n=3-5). **(B-E)** Growth curves are displayed as OD_600_ over time. The data are an average of all strains with ± SEM shaded. The dot plot shows the average AUC for each strain with a solid black line to indicate median. **(B)** Growth in human urine of clinical isolates from polymicrobial (red) vs monomicrobial (blue) UTIs. Growth in LB of clinical isolates **(C)** with *exoU* (red) versus without *exoU* (blue), **(D)** with *pilA* (red) versus without *pilA* (blue), and **(E)** with *aprA* (red) versus without *aprA* (blue). Statistical significance was determined via Mann-Whitney U test (*P<0.05, **P<0.005).

### Exotoxin genes predict hemolytic activity in clinical isolates

Iron acquisition and cytotoxicity are critical determinants of *P. aeruginosa* pathogenicity, influencing both survival in the iron-limited urinary tract and the extent of host tissue damage^1,29^. To assess these traits among our clinical isolates, we quantified iron acquisition using chrome azurol S (CAS) plates and hemolytic activity using blood agar. Most strains demonstrated either strong iron acquisition or hemolytic activity, but strain SL819 was an exception that displayed increased activity in both assays (**Fig. 4A**). Siderophore production was significantly increased in strains carrying virulence factors *aprA* (**Fig. 4B**) and *pvdA* (**Fig. 4C**), by 16.1% and 6.1%, respectively. In contrast, isolates with novel sequence types (**Fig. 4D**) showed a 6.2% decrease in iron chelation compared to known MLST strains. Hemolytic activity was enhanced by 18.4% in isolates with *exoU* (**Fig. 4E**) and 44.5% in those encoding *rhlR* (**Fig. 4F**), consistent with the known roles of these genes in blood cell lysis^44,45^. In contrast, strains resistant to levofloxacin, aztreonam, or meropenem demonstrated reduced hemolytic activity (**Fig. 4G**), suggesting a potential trade-off between cytotoxicity and antibiotic resistance. Furthermore, O6 serotype strains were associated with 6.6-fold higher blood urine concentrations than other serotypes (**Fig. S7**). Collectively, these findings highlight that serotype and toxin gene presence predict host damage and may also influence the urinary niche, while resistance traits may reduce cytotoxic ability.

**Figure 4.**
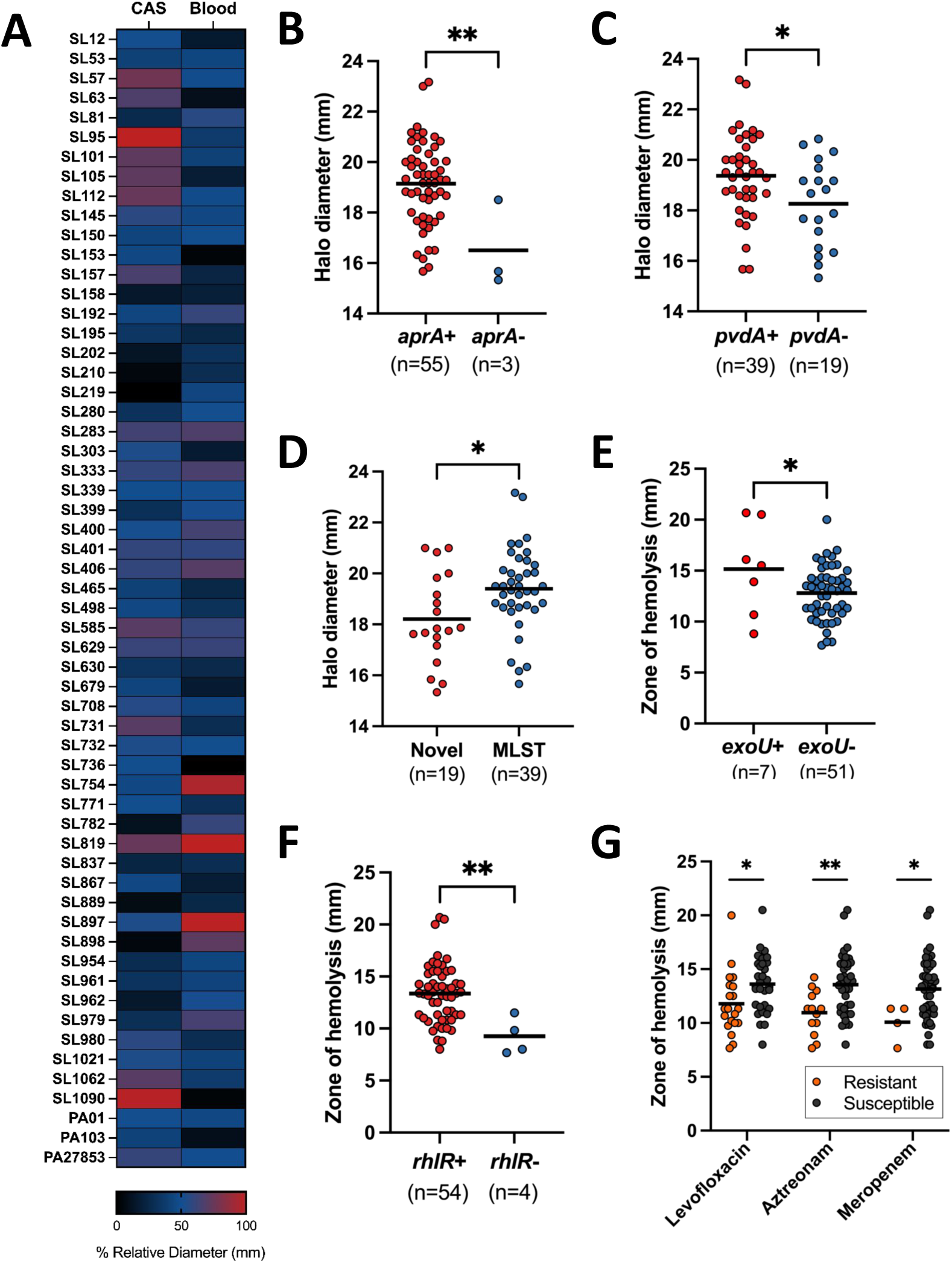
Toxin-encoding genes and antibiotic resistance influence siderophore production and hemolytic activity. *P. aeruginosa* clinical isolates and type strains were tested on chrome azurol S (CAS) plates to assess siderophore production and blood agar plates to assess hemolytic ability. **(A)** Heatmap of normalized CAS and blood agar plate zones of inhibition (mm). CAS halo diameter (mm) by presence or absence of genes **(B)** *aprA*, **(C)** *pvdA*, and by **(D)** ST type novelty. Each dot represents the average of biological replicates for an individual strain, and black lines indicate the mean of all strains in a group. Hemolytic zone of clearance (mm) in strains with and without genes **(E)** *exoU* and **(F)** *rhlR*. **(G)** Zone of hemolysis (mm) by resistance to levofloxacin, aztreonam, and meropenem. Resistant strains are shown as orange dots, and susceptible strains are black dots. Mean is indicated as a solid line. Statistical significance was determined via unpaired t-test (*P<0.05, **P<0.005).

Motility is enhanced among novel *P. aeruginosa* UTI isolates.

Motility and biofilm formation are central to *P. aeruginosa* persistence in the urinary tract, allowing tissue colonization and evasion of antibiotic clearance^1,9,46^. To evaluate these traits, we assessed swimming motility and biofilm production *in vitro* across our clinical isolates. Despite all isolates encoding the flagellar machinery, 5 isolates were non-motile, while the rest demonstrated varying levels of motility (**Fig. 5A**). Isolates from patients with recurrent urinary tract infections (rUTI) were 5.5 times more likely to carry the type A flagellin gene (*flaA*), whereas isolates from non-rUTI patients were more likely to carry type B flagellin (*fliC*) (**Fig. 5B**). Importantly, isolates belonging to novel sequence types exhibited 20.2% greater swim motility compared to those with established sequence types (**Fig. 5C**). Increased motility was associated with the presence of the quorum-sensing regulator *rhlR* (**Fig. 5D**), which also affects biofilm formation. Biofilm formation was 2.2 and 2.4 times higher in strains of the O3 serotype (**Fig. 5E**) and strains harboring the transcriptional regulator *lasR* (**Fig. 5F**), respectively. In contrast, isolates resistant to levofloxacin exhibited 51.8% weaker biofilm formation than those that were susceptible (**Fig. 5G**). These trends suggest that motility and biofilm capacity may act as complementary strategies for persistence in the urinary tract.

**Figure 5.**
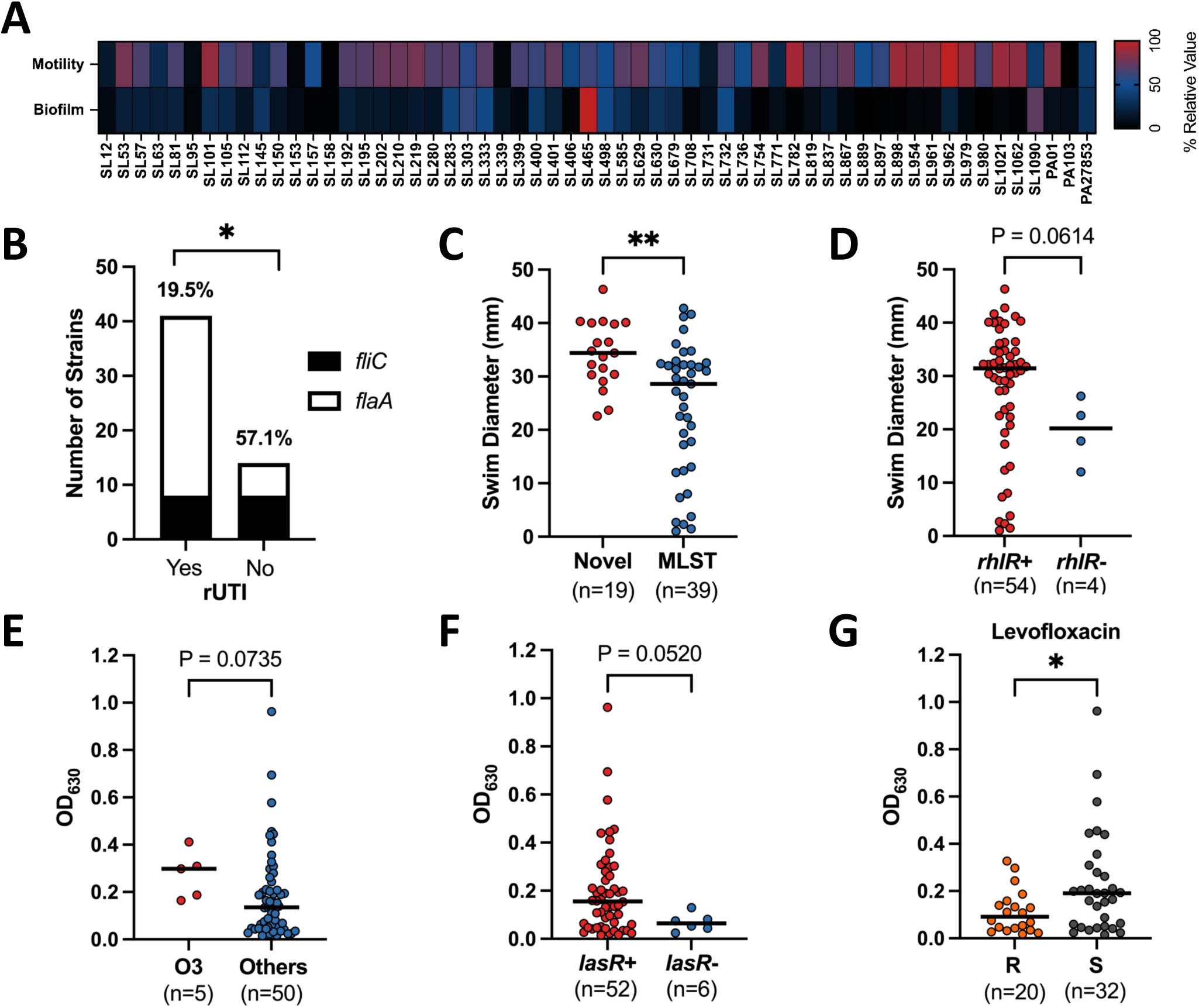
Motility and biofilm phenotypes correlate with virulence genes, Levofloxacin resistance, and recurrent UTI. **(A)** Heatmap of normalized motility swim diameter (mm) and biofilm formation in LB (optical density at 630 nm). **(B)** Prevalence of flagellum encoding gene type A (*flaA*, shown in white) and type B (*fliC*, shown in black) in strains isolated from patients with recurrent UTI (rUTI) (red) compared with patients without rUTI (blue). Motility (swim diameter) in strains with **(C)** novel ST types (red) versus those with established MLST (blue) and **(D)** isolates with the *rhlR* gene (red) compared to those without (blue). Dots represent the mean of biological replicates for a strain, and bars indicate median. Biofilm formation (OD630) in strains **(E)** of the O3 serotype (red) compared to other serotypes (blue), **(G)** with (red) versus without (blue) the *lasR* gene, and **(F)** resistant (R, orange) versus susceptible (S, black) to levofloxacin. Statistical significance was determined via Fisher’s exact test for panel B and Mann-Whitney U test for panels C-G (*P<0.05, **P<0.005).

### Swimming motility and ExoS drive colonization and persistence of uropathogenic *P. aeruginosa* in the murine model of UTI

To complement our *in vitro* assay findings, we used the traditional ascending murine UTI model to directly assess how phenotypic and genotypic characteristics contribute to colonization and persistence *in vivo*^47^. Four *P. aeruginosa* strains (PAO1, PA103, SL158, and SL192) were used to infect CBA/J mice (n=20) via transurethral inoculation.

Strains were selected to represent distinct Type III secretion system toxin profiles, enabling comparison between *exoS*-dominant and *exoU*-dominant genotypes. Urinary bacterial burden was quantified over a 96-hour period through serial CFU enumeration from urine samples, followed by organ collection and plating at the endpoint to determine bacterial load in the bladder, kidneys, and spleen. PAO1 consistently maintained the highest urine CFU burden, whereas PA103 exhibited a decline in CFU/mL over time (**Fig. 6A, Fig. S8**). A similar pattern was seen in tissue CFU burdens, where PA103 was unable to establish organ colonization, the two clinical isolates displayed intermediate levels, and PAO1 colonized very well (**Fig. 6B**). Kidney colonization differed by motility phenotype (**Fig. 6C**), with motile strains colonizing the kidney at median levels over one log higher than non-motile strains. Quantification of organ and urine CFU revealed differences in colonization capacity associated with the presence of key virulence genes (**Fig. 6D**). Strains lacking the toxin gene *exoS* exhibited a 100-fold decrease in the urine and a 5-fold decrease in both bladder and kidney colonization compared to *exoS*-positive strains (**Fig. 6D**). Additionally, isolates expressing type B flagellin (*fliC*) showed a 10.9-fold advantage in bladder colonization and a 13.5-fold advantage in kidney colonization relative to those expressing type A flagellin (*flaA*). To contextualize these *in vivo* findings, we compared the prevalence of these same virulence genes across *P. aeruginosa* genomes from other infection sources from the PATRIC database (**Fig. 6E**). We found that UTI isolates were significantly enriched for *exoS* compared to lung isolates. Collectively, these findings indicate motility and exotoxin ExoS are key fitness factors essential to *P. aeruginosa* colonization, persistence, and ascension in the urinary tract.

**Figure 6.**
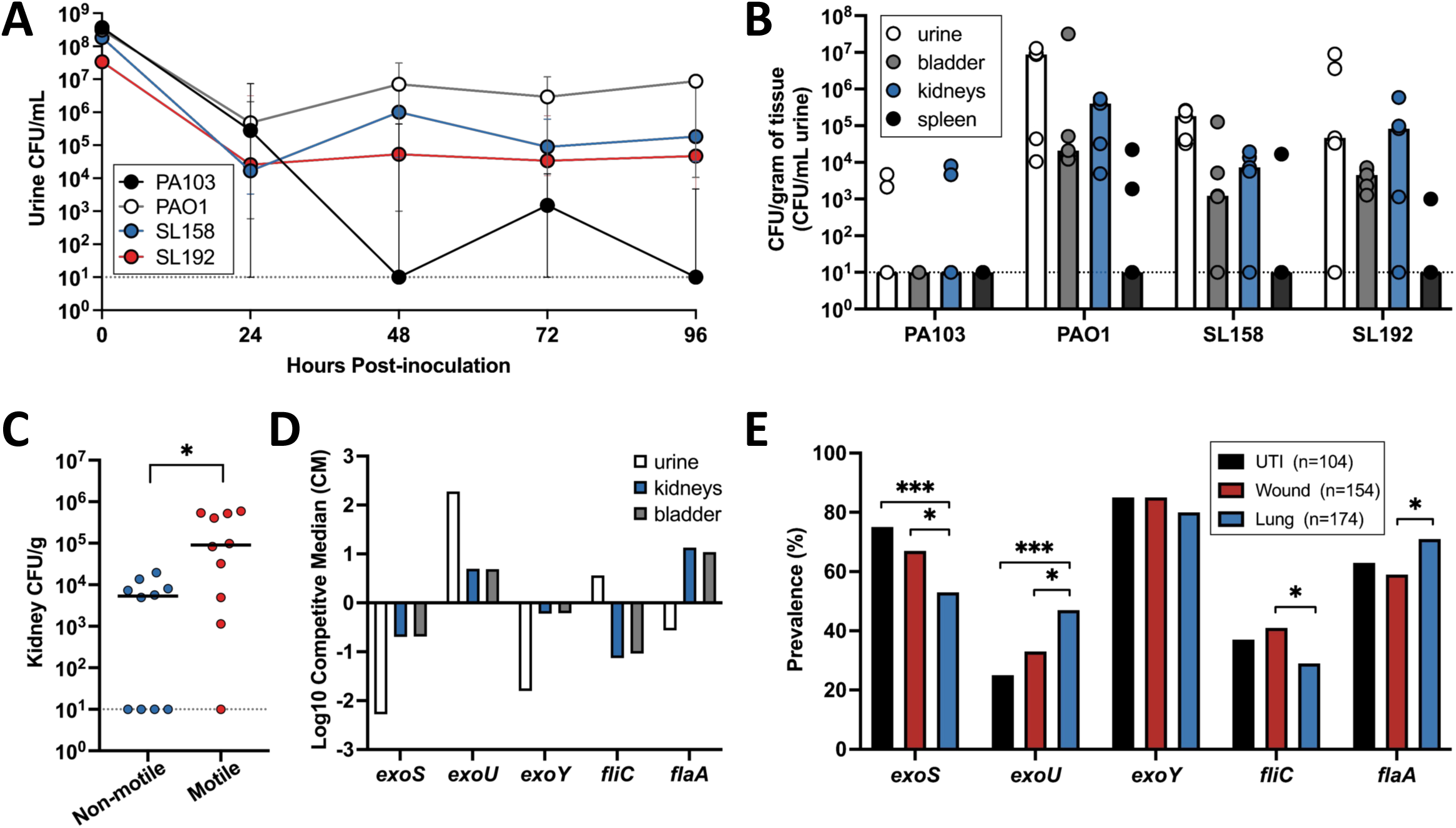
Murine UTI model links virulence traits to bacterial burden in urine and organs. Female CBA/J mice (n=20) were transurethrally inoculated with 2x10^8^ CFU of *Pseudomonas*, either PAO1 (n=5), PA103 (n=5), SL158 (n=5), or SL192 (n=5). **(A)** Urine samples were collected from each mouse every 24 hours post-infection and plated to determine CFU burden. Mean urine CFU of biological replicates (n=5) is displayed (dot) ± SEM. The limit of detection is indicated by a dashed line. Mice were sacrificed after 96 hours, and organs were aseptically collected, homogenized, and plated to determine CFU per gram of tissue. **(B)** 96-hour mean organ CFU and urine CFU/mL for each mouse (dot) are indicated (n=20) with bar height indicating median and a dashed line indicating the limit of detection. **(C)** Plot of kidney CFU burden in non-motile (PA103, SL158) compared to motile (PAO1, SL192) strains. Significance was determined by Mann-Whitney U test (*P<0.05). **(D)** Log_10_ competitive median of bladder (grey), kidney (blue), and urine (white) CFU between the presence and absence of exotoxins and flagella. **(E)** Bar graph showing the proportion of *exoY*, *exoS*, *exoU*, *fliC*, and *flaA* positive *P. aeruginosa* isolates stratified by infection type: UTI (black), wound (red), and lung (blue). Significance was determined with Fisher’s exact test (*P<0.05, ***P<0.0005).

## DISCUSSION

In this study, we performed a comprehensive genotypic and phenotypic characterization of *Pseudomonas aeruginosa* strains isolated from urinary tract infections (UTIs) within our regional healthcare system. While *P. aeruginosa* has been extensively studied in the context of respiratory and wound infections, its role in UTIs has been understudied by comparison. The absence of a dedicated UTI reference strain underscores this gap. Indeed, our experimental work relied on type strains derived from wound (PAO1), lung (PA103), and bloodstream (PA27853) infections^34–37^, demonstrating the need for dedicated UTI reference strains and models to advance understanding of *P. aeruginosa* uropathogenesis.

Our university healthcare system has a higher-than-expected prevalence of uropathogenic *P. aeruginosa*. Additionally, 34.5% of our isolates collected represented novel sequence types, indicating previously uncharacterized pathogen lineages circulating in our region. These novel isolates displayed a unique phenotypic profile including enhanced motility and reduced iron-chelating activity. Despite these shared phenotypes, phylogenetic analysis revealed high genomic diversity, with no distinct clade encompassing these novel sequence types. Resistance to the monobactam antibiotic aztreonam was also enriched among novel sequence types, proposing that emerging genotypes may be co-acquiring resistance alongside other traits relevant to urinary tract colonization. This degree of genomic novelty suggests the emergence of previously undescribed urinary-adapted lineages, potentially driven by selective pressures unique to our geographic region. Given the known heterogeneity of many *P. aeruginosa* virulence and resistance loci, these findings further support the need for a UTI-specific prototype strain that reflects the genomic features of urinary lineages.

O-antigen serotyping identified eight distinct serotypes in our cohort, with O6, O11, and O5 comprising the majority of isolates. These serotypes have also been among the most prevalent in ventilator-associated pneumonia and burn wound isolates^30,31,48^. Interestingly, we found previously undescribed correlations between serotypes and patient data in our cohort. O6 strains were more common in patients with indwelling urinary catheters (IDCs) and were also linked to elevated blood in the urine, while O4 strains were more common in patients with diabetes mellitus (DM). Together, these findings highlight the value of integrating genotypic characterization with patient data to better define patterns of *P. aeruginosa* infection across diverse patient cohorts.

Although antibiotic resistance rates in our patient population were consistent with those in healthcare-associated infections^49^, we discovered novel ties between resistance phenotypes and host demographics. Notably, isolates from African American patients exhibited significantly higher resistance to meropenem and aztreonam than those from White patients, suggesting possible population-level differences in prior exposures or environmental reservoirs. Strains of serotype O11 were more likely to be extensively drug-resistant (XDR), consistent with previous studies correlating O11 to antibiotic resistance^50^. As described in other infection models^51,52^, we observed a trade-off between biofilm formation and antibiotic resistance. Interestingly, resistance to levofloxacin, aztreonam, or meropenem was associated with reduced hemolytic activity, raising the possibility of previously undescribed fitness trade-offs between resistance and cytotoxicity in the context of uropathogenic *P. aeruginosa*.

To translate these *in vitro* phenotype findings *in vivo*, we utilized the murine model of ascending UTI. To date, there is a paucity of literature utilizing this model with *P. aeruginosa*, with most prior work focused on catheter-associated infection^53,54^. Studies in LACA mice have shown quorum sensing systems including PQS, Rhl, and Las are crucial for *P. aeruginosa* colonization and persistence in the urinary tract^13,55^. In our CBA/J model, previously only used in the context of *P. aeruginosa* co-infections^56,57^, swimming motility was a key predictor of urinary tract colonization. This pattern aligns with observations in other uropathogenic species^3,58^ and is consistent with the established role of motility in *P. aeruginosa* dissemination across multiple infection sites, including lung^46,59^, wound^60^, and burn^61^ models.

Flagellin type also influenced colonization outcomes. PAO1, which carries type B flagellin (*fliC*), exhibited a modest colonization advantage over strains encoding type A flagellin (*flaA*). Interestingly, flagellin type also correlated with recurrent UTI; *flaA* was enriched in isolates from patients with recurrent UTIs, whereas *fliC* was more common in non-recurrent cases. This may indicate selection to avoid immune recognition of FliC by TLR5^62^. Indeed, FliC is very immunogenic and has proven effective as a therapeutic agent against *P. aeruginosa* in experimental models^63–66^.

In addition to motility, we also found exotoxin profiles play a role in *P. aeruginosa* colonization. In a previous study, the type 3 secretion system (T3SS), specifically the injection of ExoU, was found to be the main contributor to virulence in acute catheter-associated urinary tract infection (CAUTI)^67^. The T3SS is also important in lung and wound pathogenesis, but there is debate over whether *exoS* or *exoU* plays a larger role^68–70^. In our murine UTI model, *exoS*-positive strains (SL158, PAO1) exhibited significantly higher bacterial burdens than *exoU*-positive strains (SL192, PA103), suggesting ExoS plays a more prominent role in urinary tract colonization. Supporting this, when analyzing urinary isolates from the PATRIC database, we found that *exoS* was enriched in urinary *P. aeruginosa* compared to lung isolates. Larger *in vivo* cohorts will be necessary for refining these conclusions and clarifying the mechanisms of *P. aeruginosa* dissemination and persistence in the urinary tract, as has been done in similar studies with *E. coli*^71^.

While many previous studies have evaluated PAO1 in the murine UTI model, to our knowledge, none have assessed PA103. Notably, PA103 was the only strain unable to colonize the murine urinary tract, posing the question of what phenotypic and genotypic factors are contributing to this defect. PA103 is non-motile due to a single amino acid substitution in *fleQ*^72^, a mutation absent from the other three strains tested *in vivo*. In fact, none of our clinical UTI strains exhibited this mutation, further demonstrating motility is essential to *P. aeruginosa* urovirulence. PA103 also possesses a *lasR* loss-of-function (LOF) mutation that impairs quorum sensing and protease expression, a hallmark commonly seen in chronic *P. aeruginosa* infections such as cystic fibrosis^73–75^. In our UTI isolates, *lasR*-deficient strains demonstrated reduced biofilm formation, consistent with the role of *lasR* in regulating quorum sensing-associated virulence pathways^10^. The Las system has indeed been proven crucial to *P. aeruginosa* pathogenesis in the urinary tract^14^. Future murine studies restoring motility in PA103 via *fleQ* repair will provide critical insight into the relative contributions of lost motility versus impaired quorum sensing to its colonization defect.

Overall, our data reveal that *P. aeruginosa* UTIs in our region are caused by a genetically diverse set of isolates, many of which represent novel lineages with unique combinations of virulence and resistance traits. By integrating genomic, phenotypic, and *in vivo* analyses, we provide new insights into how *P. aeruginosa* adapts to and persists within the urinary tract. These findings highlight the need to better recognize *P. aeruginosa* as a clinically significant uropathogen and to invest in research that addresses this underexplored but clinically important role.

## MATERIALS AND METHODS

### Bacterial Strains, Media, and Culture Conditions

Bacterial strains used in this study included clinical isolates of *Pseudomonas aeruginosa* collected under University of South Alabama (USA) IRB protocol #2178590. All clinical isolates were obtained from the University Hospital from the urine culture. Strains were confirmed to be *Pseudomonas aeruginosa* via an oxidase test and/or Matrix-Assisted Laser Desorption/Ionization Time-Of-Flight (MALDI-TOF). Isolates were coded and stored at -80°C in Luria Broth (LB) containing 20% glycerol. Strains were cultured from a single colony in LB, which contains 0.5 g NaCl, 5 g yeast extract, and 10 g tryptone per liter. To prepare LB agar plates, 5 g of agar was added. Cultures were incubated at 37°C with aeration at 200 rpm unless otherwise stated. In experiments requiring urine, filter-sterilized (0.22 µm) pooled human urine from at least six healthy de-identified female donors was used (deemed IRB exempt by USA).

### Antibiotic Susceptibility Testing, Urinalysis, and Patient Metadata

Antimicrobial susceptibility testing (AST) and urinalysis were performed at the University of South Alabama hospital microbiology lab. AST was performed using an automated broth dilution protocol (BD Phoenix™ M50) according to Clinical and Laboratory Standards Institute (CLSI) guidelines. Strains were classified as resistant (R), intermediate (I), or susceptible (S) based on minimum inhibitory concentration (MIC). Antibiotic-specific MIC ranges used to make R, I, or S determinations are included in **Supplemental Data File 1**. Urinalysis was performed via dipstick chemical analysis and microscopy.

Urine culture, AST, and urinalysis results associated with the clinical isolates were obtained via retrospective chart review (IRB protocol #2178590). From these reports, causative agents, antibiotic susceptibilities, and dipstick results were recorded. In addition, de-identified patient variables including patient age, race, sex, and body mass index (BMI) were collected from the patient charts and associated with the coded isolate.

### Biofilm Assay

Biofilm formation was assessed using a crystal violet staining assay. Overnight cultures of strains were diluted 1:100 into LB or filter-sterilized human urine in duplicate in 12-well plates. The plate was sealed with a gas-permeable membrane and incubated statically at 37°C for 24 hours. Wells were washed with 1X phosphate-buffered saline (PBS) and stained with 0.1% crystal violet for 10 minutes. The stain was aspirated, and the wells were washed a second time. Biofilm biomass was quantified by solubilizing the dye with ethanol and measuring absorbance via optical density at 630 nm (OD_630_) using an Agilent BioTek 800 TS Absorbance Reader.

### Iron Acquisition, Hemolysis, and Motility Assays

Iron chelation and siderophore production were measured using the Chrome Azurol S (CAS) assay. CAS agar was prepared as previously described^76^. For each strain, 5 µL of overnight culture was spotted onto a CAS agar plate and incubated for 16 hours at 37°C. The halo diameter was then measured and recorded in millimeters (mm). Hemolytic activity was evaluated by spotting 5 µL overnight bacterial cultures onto 5% Sheep Blood in Tryptic Soy Agar plates (Hardy Diagnostics CAT# A10). Plates were incubated at 37°C for 16 hours, then zones of clearance were measured in mm.

To assess motility, strains were tested in semi-soft agar. Overnight cultures of each strain were normalized to an OD_600_ of 10.0, then resuspended in HEPES buffer (pH 8.4). With an inoculating needle, cultures were stabbed into tryptone agar plates with the following composition per liter: 10 g tryptone, 5 g sodium chloride, and 2.5 g agar. Plates were incubated for 16 hours at 30°C, then the swimming diameter was measured in mm.

### Growth Curve Assay

Overnight cultures were washed once in 1X PBS, then diluted 1:100 into LB or filter-sterilized, pooled human urine in a 96-well plate. The plate was sealed with a gas-permeable membrane, and bacterial growth was monitored by measuring OD₆₀₀ over time using a BioTek LogPhase600 Microbiology Reader. Readings were taken every 10 minutes for 24 hours.

### Murine Model of UTI

Female CBA/J mice between the ages of 6 to 8 weeks old were acquired from Jackson Laboratories. Under ketamine/xylazine anesthesia, mice were transurethrally inoculated with 50 µL of 2 x 10^8^ CFU/mL bacterial suspension of strain PAO1, PA103, SL158, or SL192 with a sterile polyethylene catheter connected to an infusion pump^47,77^. Urine samples were collected and plated on LB agar every 24 hours post-inoculation to assess bacterial load. After 96 hours, the mice were euthanized, and the bladders, kidneys, and spleens were aseptically removed and homogenized. Homogenates were then serially diluted and plated on LB agar to assess bacterial burden. All protocols were approved by the Institutional Animal Care and Use Committee (IACUC #2006187) at the University of South Alabama.

### Whole-Genome Sequencing

Genomic DNA was extracted using the Promega Wizard^®^ Genomic DNA Purification Kit following the manufacturer’s instructions. Libraries were prepared using standard Illumina protocols to produce paired end 150 bp reads. Isolates SL12-SL736 were sequenced by Moderna TX via the NovaSeq 6000 system, and isolates SL754-SL1090 were sequenced by SeqCoast Genomics using the Illumina NextSeq 2000 platform. Raw sequencing data has been deposited in SRA (PRJNA1390275).

### Bioinformatics Analysis

Sequencing reads were processed using FastQC^78^, and genome assemblies were generated with Unicycler^79^ on BVBRC using an annotated recipe for the *P. aeruginosa* taxonomy. Eight genomes were flagged as poor quality after this process (SL53, SL157, SL158, SL192, SL210, SL219, SL280, and SL339) and were subjected to the following polishing process. Reads were trimmed with fastp^80^, and, where possible, taxonomically filtered with Kraken2^81^ to retain *Pseudomonas* (taxid 286). Genomes were assembled using SPAdes^82^, polished once with Pilon^83^ using bwa-mem2 read-mapping. Contigs <1 kb were discarded, and assembly metrics were assessed with QUAST^84^. Once all assemblies were checked for quality, annotations were performed using the RAST tool kit (RASTtk)^85^. Assembly quality metrics for all strains are included in **Supplemental Data File 2**.

SNP analysis was performed via kSNP4 to construct a maximum likelihood phylogenetic tree, and iTOL^86^ was utilized for editing the tree. Gene presence was determined by BLASTP (see **Supplemental Data File 3** for sequences used) with a threshold of >85% identity and >90% query cover. ST typing was performed using the PubMLST *Pseudomonas aeruginosa* typing database^39^. Allelic matches are provided in Table 2. Serotypes were determined using the PAst v1 *in silico* serotyping tool^40^. To determine the presence of *fleQ* or *lasR* mutations, multiple sequence alignment was performed via Mafft. *P. aeruginosa* isolates from the PATRIC database stratified by infection type (UTI, wound, and lung) genome accession number can be found in **Supplemental Data File 4**.

### Statistical Analysis

Competitive median (CM) was computed as 1/(median CFU of isolates with the gene of interest)/(median CFU of isolates without the gene of interest). Statistical analyses were performed using GraphPad Prism (v10.4). Variables were screened for correlation via Pearson’s correlation. Shapiro-Wilk normality tests and D’Agostino & Pearson omnibus normality tests were used to determine normality of data, and variances were compared using the F test. Group comparisons were made using unpaired two-tailed t-tests or Mann-Whitney U tests (95% CI) based on data distribution. When comparing two categorical variables, Fisher’s exact test was used. P values < 0.05 were considered statistically significant.

### Ethics

This study was approved by the University of South Alabama Institutional Review Board (IRB, protocol #2178590). Human derived bacterial strains were obtained in accordance with the approved protocol. All bacterial isolates were coded prior to entering the research laboratory to ensure patient confidentiality. All animal protocols were approved by the Institutional Animal Care and Use Committee (IACUC, protocol #2006187) at the University of South Alabama College of Medicine.

## ACKNOWLEDGMENTS

We would like to thank the University of South Alabama Frederick P. Whiddon College of Medicine for the start-up funds to support this project in the laboratory of Dr. Allyson Shea. We would also like to thank Teresa Barnett for the collection of bacterial isolates.

## SUPPLEMENTAL MATERIAL

**Supplemental Figure 1. UTI pathogen distribution in Mobile, Alabama.** Retrospective chart review was conducted to identify the top UTI causative agents across the USA healthcare system from June 1, 2024 to May 31, 2025. Positive UTI cases were identified via urine culture (n=6929). Organisms <1% are not shown.

**Supplemental Figure 2. O11 serotype is correlated with extensive drug-resistance (XDR).** Prevalence of extensive drug-resistance (XDR, shown in black) in strains with the O11 serotype versus all other serotypes. Fisher’s exact test was used to determine statistical significance (*P<0.05).

**Supplemental Figure 3. Clinical P. aerugi*nosa* strains show a high number of antibiotic resistance genes.** The genomes of the 55 clinical *P. aeruginosa* isolates were blasted for antibiotic resistance genes in the Comprehensive Antibiotic Resistance Database (CARD). Perfect hits (dark blue) have all amino acids matching the CARD sequence, strict (light blue) hits indicate very close matches with slight variations in the sequence. White indicates the gene is absent.

**Supplemental Figure 4. Correlation matrix of genotypic data to patient variables.** Gene presence or absence was determined by BLAST and correlated to patient variables. Gene presence or absence and patient variables were tested for correlation in GraphPad Prism with Spearman r. Red indicates a positive correlation, and blue indicates a negative correlation. P values were determined with 95% confidence (*P<0.05, **P<0.005, ***P<0.0005, ****P<0.00005). Blank or excluded comparisons are indicated with an X through the cell.

**Supplemental Figure 5. Growth in human urine is affected by virulence gene presence and O-antigen serotype. (A)** Dot plot with growth in human urine in strains with (red) and without (blue) genes *parS*, *armR*, and *toxA*. Bar height indicates median AUC. **(B-C)** Plots of growth in human urine (AUC) in serotypes **(B)** O5 and **(C)** O11 (red) versus all other serotypes (blue). Each dot represents a clinical isolate. The median of each group is indicated with a black line. Statistical significance was determined via Mann-Whitney U test (*P<0.05).

**Supplemental Figure 6. Correlation matrix of phenotypic data to patient variables.** The 55 *P. aeruginosa* clinical isolates were subjected to Chrome Azurol S (CAS), blood agar, biofilm, and growth curve assays to obtain phenotypic data. Patient variables were obtained and tested for correlation with phenotypic data in GraphPad Prism with Spearman r. Red indicates a positive correlation, and blue indicates a negative correlation. P values were determined with 95% confidence (*P<0.05, **P<0.005, ***P<0.0005).

**Supplemental Figure 7. Patients with strains of O6 serotype have higher levels of blood in urine.** Urinalysis results from USA Hospital were obtained for the UTI urine samples of origin for the 55 clinical isolates. A plot is shown of blood cells in urine specimens containing *P. aeruginosa* strains of serotype O6 versus other serotypes. Statistical significance was determined via Mann-Whitney U test (*P<0.05).

**Supplemental Figure 8. Murine urine CFU burden of *P. aeruginosa* clinical and type strains.** Urine was plated every 24 hours to assess bacterial load. Colored symbols indicate the mean of three technical replicates for each individual mouse (n=5).

**Supplemental Data File 1:** Minimum inhibitory concentration (MIC) ranges used for antimicrobial resistance determinations.

**Supplemental Data File 2:** Assembly quality metrics for the 55 *P. aeruginosa* urinary isolates.

**Supplemental Data File 3:** Protein sequences used for BLAST analyses.

**Supplemental Data File 4:** Strains from the PATRIC database used for comparison across isolation sources.

